# Preparing for a second attack: a lesion simulation study on network resilience after stroke

**DOI:** 10.1101/2021.09.21.461167

**Authors:** Mitsouko van Assche, Julian Klug, Elisabeth Dirren, Jonas Richiardi, Emmanuel Carrera

## Abstract

**Background and Purpose:** Does the brain become more resilient after a first stroke to reduce the consequences of a new lesion? Although recurrent strokes are a major clinical issue, whether and how the brain prepares for a second attack is unknown. This is due to the difficulties to obtain an appropriate dataset of stroke patients with comparable lesions, imaged at the same interval after onset. Furthermore, timing of the recurrent event remains unpredictable.

**Methods:** Here we used a novel clinical lesion simulation approach to test the hypothesis that resilience in brain networks increases during stroke recovery. 16 patients with a lesion restricted to the primary motor cortex were recruited. At 3 time points of the index event (10 days, 3 weeks, 3 months), we mimicked recurrent infarcts by deletion of nodes in brain networks (resting-state fMRI). Graph measures were applied to determine resilience (global efficiency) and wiring cost (mean degree) of the network.

**Results:** At 10 days and 3 weeks after stroke, resilience was similar in patients and controls. However, at 3 months, while motor function had fully recovered, resilience to clinically representative simulated lesions was higher compared to controls (cortical lesion p=0.012; subcortical: p=0.009; cortico-subcortical: p=0.009). Similar results were found after random (p=0.012) and targeted (p=0.015) attacks.

**Conclusion:** Our results suggest that, after a lesion, brain networks reconfigure to increase resilience to future insults. Lesion simulation is an innovative approach, which may have major implications for stroke therapy. Individualized neuromodulation strategies could be developed to foster resilient network reconfigurations after a first stroke to limit the consequences of future attacks.

## INTRODUCTION

Does the brain become more resilient to further events, following a first stroke? Despite major progress in secondary prevention, recurrent strokes are frequent, occurring in up to 20% of patients within 3 months of onset^1^. In stroke animal models, the behavioral impact of a second lesion decreases with time^2^, suggesting that resilience to new events - defined here as the capacity of the brain to resist, overcome, or thrive in the face of adversity^3^ - progressively builds up during recovery. In humans, it is still debated whether the occurrence of an ischemic event limits the consequences of a second event^4-6^. If proven to be true, the hypothesis that resilience increases within days or months after stroke to circumvent the impact of a potential recurrent stroke, may lead to major physiological and clinical implications^3^. Understanding how and why the brain reorganizes in a certain configuration after stroke could help promote neuromodulatory therapeutic strategies that will aim not only at restoring function, but also at promoting network configuration that minimize the effects of a potential recurrent stroke. Investigating resilience after a first stroke in humans is however challenging. This is due to the difficulties to obtain an appropriate dataset of patients with comparable lesions, imaged at the same interval after stroke onset. Furthermore, timing and localization of the recurrent event remain unpredictable in a given patient.

To circumvent our inability to predict a new event, we evaluated resilience in brain networks by simulating “recurrent” lesions through node deletions in a population of patients with similar infarcts restricted to the primary motor cortex. In previous studies, node deletion has been key to determine that brain networks of healthy subjects are organized to optimize the balance between integration and segregation^7-10^. By deleting nodes randomly or according to their importance in the network, it was also possible to demonstrate that brain networks architecture confers robustness despite vulnerability of central nodes^11, 12^. Deletion of contiguous nodes was only recently considered as a method to represent strokes with their anatomical characteristics, in term of size and location^13, 14^. In humans, this method seems a particularly promising alternative to empirical studies to study resilience after stroke, given the unpredictable nature of the second clinical event.

Here, we tested the hypothesis that resilience in brain networks increases during stroke recovery. For that purpose, we considered resilience as the network capacity to maintain information capacity based on the measure of the global efficiency (E_glob_). Resilience was investigated by simulating two types of attacks. In one classical approach, nodes of whole brain networks were serially deleted, randomly or based on their importance in the network. We then simulated clinically representative lesions and evaluated their impact on network reorganization. Operationally, we used lesion simulation in a population of stroke patients with a lesion restricted to the primary motor cortex and contralateral hand paresis at three time points within 3 months of onset. All patients had a detailed motor examination and functional connectivity analyses (resting-state fMRI and graph measures) at each time point.

## MATERIAL AND METHODS

### Participants

We included 16 consecutive stroke patients (6 women; age 73 ± 12 y) with a small lesion restricted to the primary motor cortex (M1) and isolated contralateral hand paresis. These patients were prospectively recruited out of the 1656 patients admitted in our stroke center during the study period. Exclusion criteria were i) left handedness, ii) significant carotid or intracranial artery stenosis (>50%), iii) history of stroke or psychiatric disease. 16 healthy subjects, matched for age, gender, and cardiovascular risk factors (6 women; 70 ± 10 yo) were included This cohort was used in a recent study with the distinct aim of investigating surrogates of motor recovery focusing on the periinfarct within three weeks after stroke.^15^ This previous study did not include graph analysis in whole brain networks, nor lesion simulation. Detailed measures of motor function and imaging data were obtained on the same day at three time points (TP) in patients: TP1; <10 days; TP2; 3 weeks and TP3; 3 months post stroke and at one time points in healthy subjects. Consent was obtained according to the Declaration of Helsinki. The study was approved by the Geneva Ethical Committee.

### Behavior Assessment

Hand motor function was evaluated by measuring hand dexterity (nine-hole pegboard task) and isometric grip strength (JAMAR dynamometer, Asimow Engineering Co., Los Angeles, CA). A two-point discrimination task applied to the index fingers was used to exclude sensory deficits. For subsequent analysis of dexterity and grip strength, performance of the paretic hand was normalized by the one of the non-paretic hand (paretic hand/unaffected hand). Owing to the non-normality of the data, Wilcoxon tests were used to examine changes in hand motor function.

### Imaging acquisition

Images were acquired on a 3T MRI (MAGNETOM Prisma, Siemens Healthcare, Erlangen, Germany; 64-channel head-coil) the same day as behavioral testing. Acquisition of resting state functional images was performed using a gradient EPI sequence (TE/TR = 30/1200 ms, voxel size = 3 mm isotropic, 400 volumes, total acquisition time 8 minutes). Continuous eye-tracking was used to check wakefulness. Respiratory movements were recorded using a transducer at the level of maximum respiratory expansion (BioPac Inc, Santa Barbara, USA). T1-weighted anatomical scans were acquired with an MPRAGE sequence (TE/TR = 2.27/2300 ms, voxel size = 1.0 mm isotropic), together with T2-weighted (TE/TR = 108/6090 ms, voxel size = 0.4 × 0.4 × 4.0 mm) and DWI images (TE/TR: 52/4300 ms, voxel size = 1.4 × 1.4 × 4.0 mm). Finally, the protocol included Brain MR angiography (TOF) and precerebral Doppler ultrasound to rule out intracranial or precerebral stenosis.

### Imaging data analysis

#### Imaging data preprocessing

Data were preprocessed using SPM12 and in-house MATLAB scripts according to an established pipeline (https://miplab.epfl.ch/index.php/software/wFC) with additional signal cleaning.^16^ First, functional images were realigned for each subject. Then, anatomical T1 images were co-registered to the mean functional image of the corresponding subject and segmented into grey matter, white matter, and cerebrospinal fluid maps. We used the Brainnetome atlas, which provides a parcellation of the human brain in 246 regions and includes a fine-grained parcellation of the motor cortex, to atlas the grey matter of each subject in native space.^17^ The resulting map was co-registered to the mean image of the functional data of the corresponding participant.

#### Extraction of brain signals

The first 5 volumes were discarded to account for magnetization equilibrium. Time-courses were linearly detrended at each voxel, averaged for each region of the atlas, and scaled by the signal mean of the given region. The six motion parameters, their first derivatives, and the average signal of CSF were regressed out. Additionally, respiratory movements were corrected using RETROICOR. To correct for remaining outlying spikes, time courses were winsorized to the 5^th^ and 95^th^ percentiles. They were then filtered into 4 frequency subbands using a wavelet transform (cubic orthogonal B-spline). We focus here on scale 4 of this decomposition (frequency range 0.03 < *f* < 0.06 Hz). We further checked for undesirable motion effects by computing the mean framewise displacements for all subjects and time points.^18^ There was no difference in motion across time points (Friedman test; χ^2^(2) = 0.462, p = 0.794) and no volumes were removed. Connectivity matrices were derived by computing pairwise Pearson correlation coefficients between the 246 regions of the Brainnetome atlas. Six regions of interest (ROIs), for which signal drop-out was observed in at least one subject, were removed, yielding a total of 240 ROIs. Finally, we flipped left and right hemisphere ROIs data within the connectivity matrices level for patients with right lesions (N=4)

#### Graph construction

Graphs were constructed following four steps. First, each connectivity matrix was normalized by its total connectivity strength and this full graph was used to calculate global connectivity strength. In the next step, each connectivity matrix was thresholded using an absolute threshold *w* > 0 to remove negative weights (**Figure 1**). A proportional (edge density) thresholding *t* was then applied, from *t* = 0 (no connection) to *t* = 1 (all connections retained) with a density increment of 0.1. This procedure allows filtering connectivity weights according to the strongest weights in a cumulative manner. Thus, this approach precludes the use of an arbitrary threshold and allows examining graph properties over a range of edge density values instead. Finally, each matrix was binarized before computing all other graph metrics. To derive efficiency and cost in the network, we used the brain connectivity toolbox.^19^

**Figure 1.**
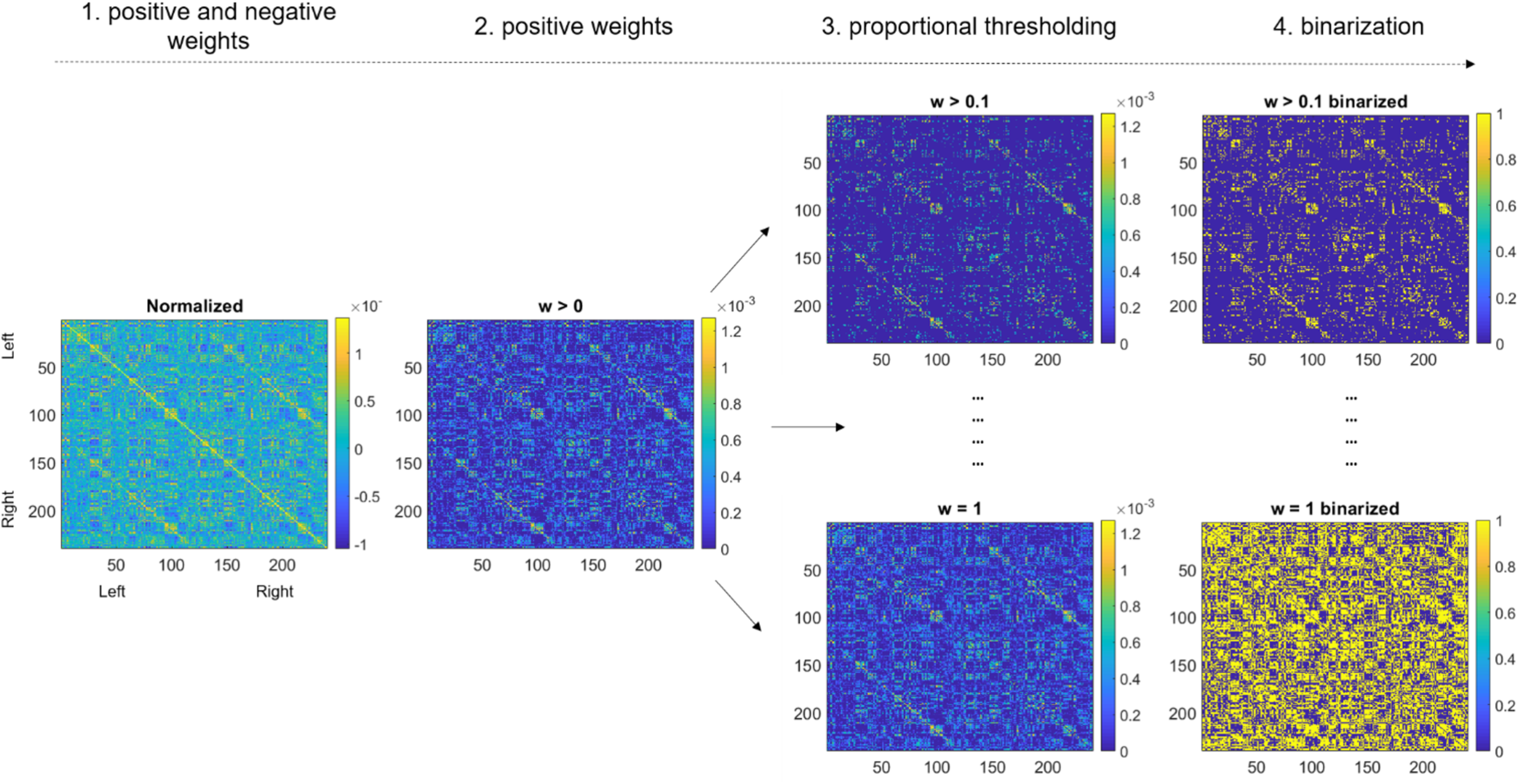
Pipeline of graph construction. 1) Fully connected graph containing positive and negative connectivity weights. 2) Application of an absolute thresholding w>0 to retain positive weights only. 3) Sparsely to densely connected graphs are obtained by means of proportional thresholding by step of incremental steps of 0.1. 4) Binarization of connectivity matrices leads to unweighted graphs.

#### Graph Metrics

The *resilience* of the network was estimated using the measure of global efficiency (*E*_*glob*_).^8, 11, 20^ This metric provides a measure of information transfer across all nodes of the network. It quantifies the extent to which nodes communicate with distant nodes. It is proportional to the inverse of the shortest path length.^21^

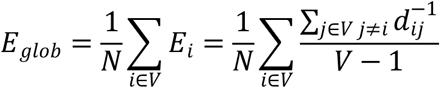

where E_i_ is the efficiency of node i, V is the set of all nodes in the network, and N is the number of nodes. (i, j) is a link between nodes i and j, (i, j ∈ V), d_ij_ is the shortest path length (distance) between i and j.

As global efficiency depends on network density, we checked the density range in which graphs remained connected in each subject. As a result, we retained the density range 0.3-1 for subsequent analyses rather than selecting arbitrary thresholds. We then derived the area under the curve (AUC) over the selected density range for E_glob_ :

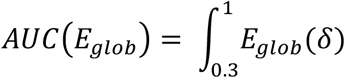

The *total wiring cost* or density of the network was estimated using the mean node degree of the network which can be estimated as the number of edges connected to each node, averaged over all nodes of the network. AUC(_Mean degree_) was computed in the same fashion as AUC(_Eglob_)

For better readability of the manuscript, we will use the terms “E_glob_ ^”^ and “mean degree” instead of “AUC(_Eglob_)” and “AUC(_Mean degree_)” in the next sections.

### Statistical Analysis

#### General Strategy

To investigate changes in resilience after stroke, we compared global efficiency (E_glob_) in whole-brain functional networks at three time points within 3 months of stroke in 16 patients with a lesion restricted to the primary motor network. We evaluated the impact of different simulated lesions (node deletion), beginning with random and targeted attacks in the whole brain network and then using attacks mimicking cortical and subcortical strokes that can be observed in clinical practice. For both the spontaneous evolution and the impact of simulated attack, we first used a linear mixed model to capture the global evolution along timepoints. T-tests were then used to compare measurements between individual timepoints and between patients and controls. Finally, we evaluated whether changes in resilience were correlated with changes in total wiring costs of the network.

##### 1. Spontaneous changes in resilience during recovery

E_glob_ was measured at each of the three time points in patients (10 days, 3 weeks, 3 months) and in controls. To evaluate changes in resilience over time in patients, we first used a linear mixed model with E_glob_ as the dependant variable, “timepoint” as a fixed effect factor and “subjects” as a random effect. Significance was evaluated by using the Satterthwaite approximations for degrees of freedom.^22^ Within patients, longitudinal comparisons between E_glob_ at the three time points were then performed using paired t-tests with false discovery rate (FDR) corrections (Benjamini–Hochberg procedure) for multiple comparisons. Between patients and controls, comparisons between the E_glob_ for patients at each time point and the E_glob_ for controls at their single time point was performed by a t-test, with FDR correction.

##### 2. Impact of lesion simulation on resilience in brain networks

###### 1) Random and targeted attacks

We first deleted nodes in random order. Global efficiency was recalculated after each node deletion (i.e. 240 times) and then averaged at each density threshold (30-100%) for each patient. AUC were then derived as described above, resulting in one value per patient at each timepoint and in one single value per control. We then performed a similar analysis with nodes deleted based on their importance within the network (targeted attack, i.e., by decreasing order of node degree). For comparison between timepoints in patients and between patients and healthy controls, we first used a linear mixed model using the same fixed and random effects as described above followed by t-tests with FDR correction for multiple comparisons.

###### 2) Clinically representative attacks

We mimicked 3 representative strokes (cortico-subcortical, subcortical, and cortical lesions (**Figure 2**)) using lesion masks corresponding to 3 patients admitted to our stroke center for infarction in the territory of the middle cerebral artery. The 3 lesion masks were manually outlined on the T2 MRI and the resulting masks normalized to MNI space with the Clinical Toolbox.^23^ Deleted nodes were defined by the intersection between the lesion masks and the ROIs from the Brainnetome atlas. As a result, the subcortical and cortical masks included 13 nodes each (respective volumes: 1.51 cm^3^ and 2.98 cm^3^) and the cortico-subcortical mask comprised 54 nodes (volume: 5.7 cm^3^). E_glob_ was computed after node deletion and the AUC was computed over the density spectrum as described above. For comparison between timepoints in patients and between patients at each timepoint we first used a linear mixed model using “timepoints” as a fixed effect, “type of lesion” and “subjects” as random effects. We then performed t-tests to compare AUC between timepoints and with healthy controls using FDR corrections for multiple comparisons as described above.

**Figure 2.**
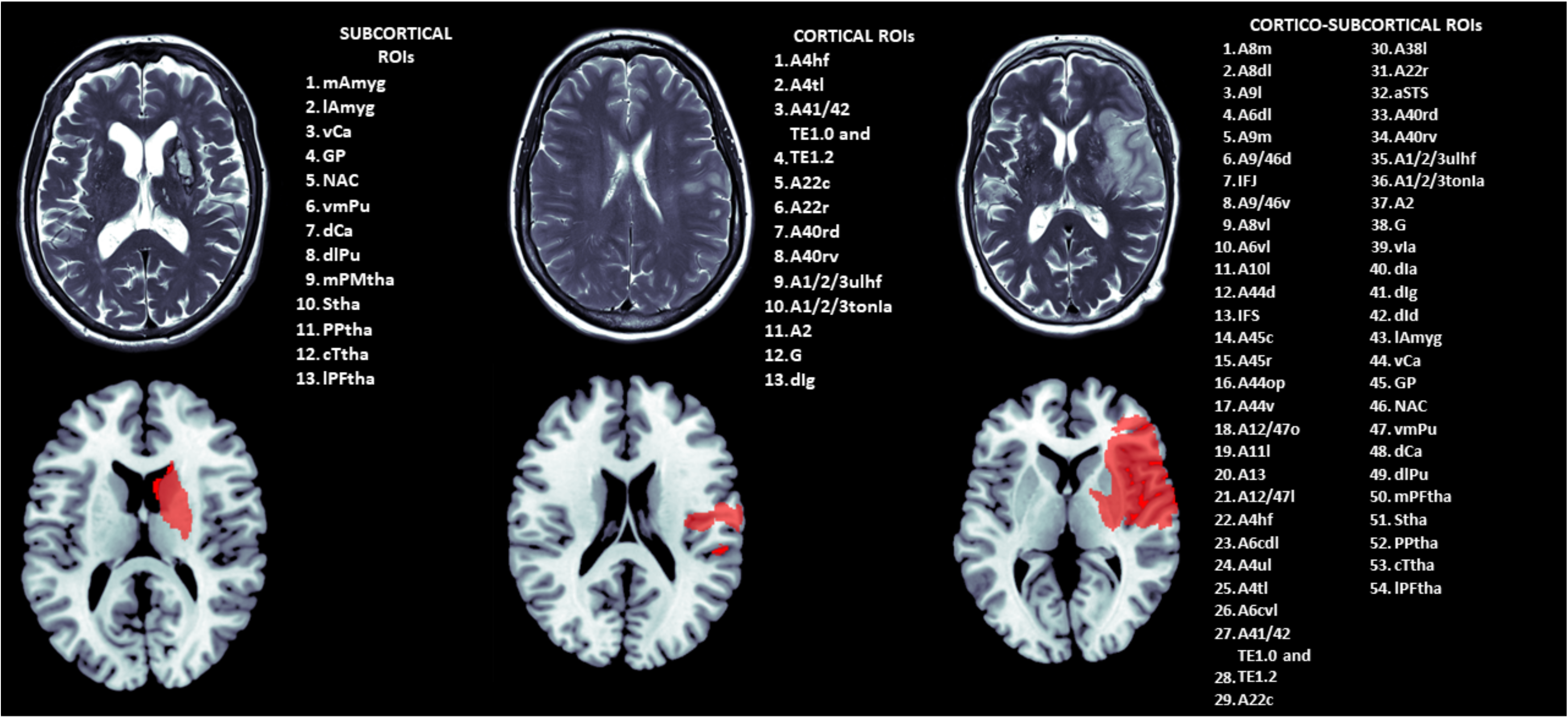
Focal Middle Cerebral Artery (MCA) strokes used for lesion simulation. Top: MRI (T2 sequence) shows the lesion for each stroke subtype (subcortical, cortical, cortico-subcortical MCA territory). Bottom: The masks (in red) correspond to normalized lesions overlaid on a standard MNI template. The list of ROIs enumerates the Brainnetome atlas regions overlapping with the lesion masks.

##### 3. Correlation between wiring cost of resilience of the networks

To determine the “price” of changes in resilience between timepoints 2 and 3, we correlated the changes in resilience (E_glob_) and the changes in total wiring cost (mean degree).

### Data availability

Anonymized data are available on request.

## RESULTS

### Changes in motor behaviour

Hand dexterity improved from TP1 to TP2 (Nine hole peg test; median laterality ratio at TP1: 1.18 (IQR: 1.06-1.71); at TP2: 1.03 (IQR: 0.94-1.27) p=0.01). At TP2, patients had fully recovered with no differences with healthy controls (median laterality ratio in controls=1.06 (IQR: 0.98-1.1; p= 0.98)). There was no change in hand dexterity between TP2 and TP3 (p=1.0). Grip strength remained stable over time (JAMAR dynamometer; TP1 to TP2: p = 0.41; TP2 to TP3: p = 0.06) and was not different from controls at any time point (TP1: p = 0.14; TP2: p = 0.21; TP3: p = 0.38).

#### 1. Changes in network resilience over time

There was a statistically significant difference between time points (mixed model, F = 5.633; p=0.009). E_glob_ was similar between TP1 to TP2 (pBH = 0.941) and between patients and controls at TP1 (pBH = 0.222) and TP2 (pBH = 0.222). However, E_glob_ increased from TP2 to TP3 (pBH = 0.017) and was higher in patients at TP3 compared to controls (pBH = 0.006). **(Figures 3**,**4)**

**Figure 3.**
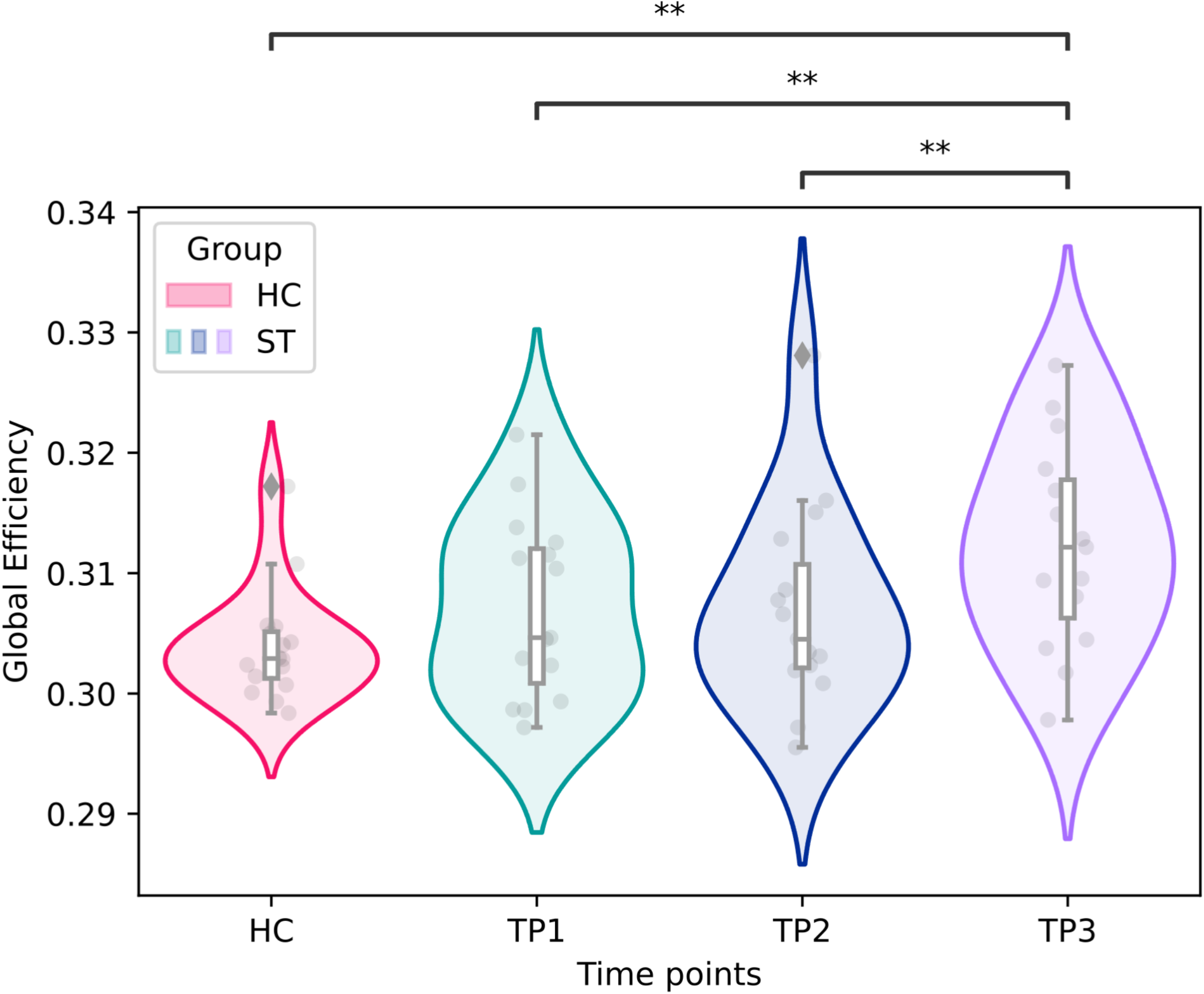
Changes in network efficiency over time. Violinplots with inner boxplots for AUCs of global efficiency in patients (ST, shown in red) and healthy controls (HC, shown in green, blue and purple) at each timepoint. Each box extends from the 25th percentile to the 75th percentile with a line indicating the median. Upper and lower whiskers show the range up to the upper and lower extremes (±1.5*inter-quartile range). Individual values are represented by grey dots. Outliers are represented by grey diamond shapes. Significance was evaluated with multiple t-tests with false discovery rate corrections; p-values are represented as * for ≤ 0.05 and ** ≤ 0.001.

**Figure 4.**
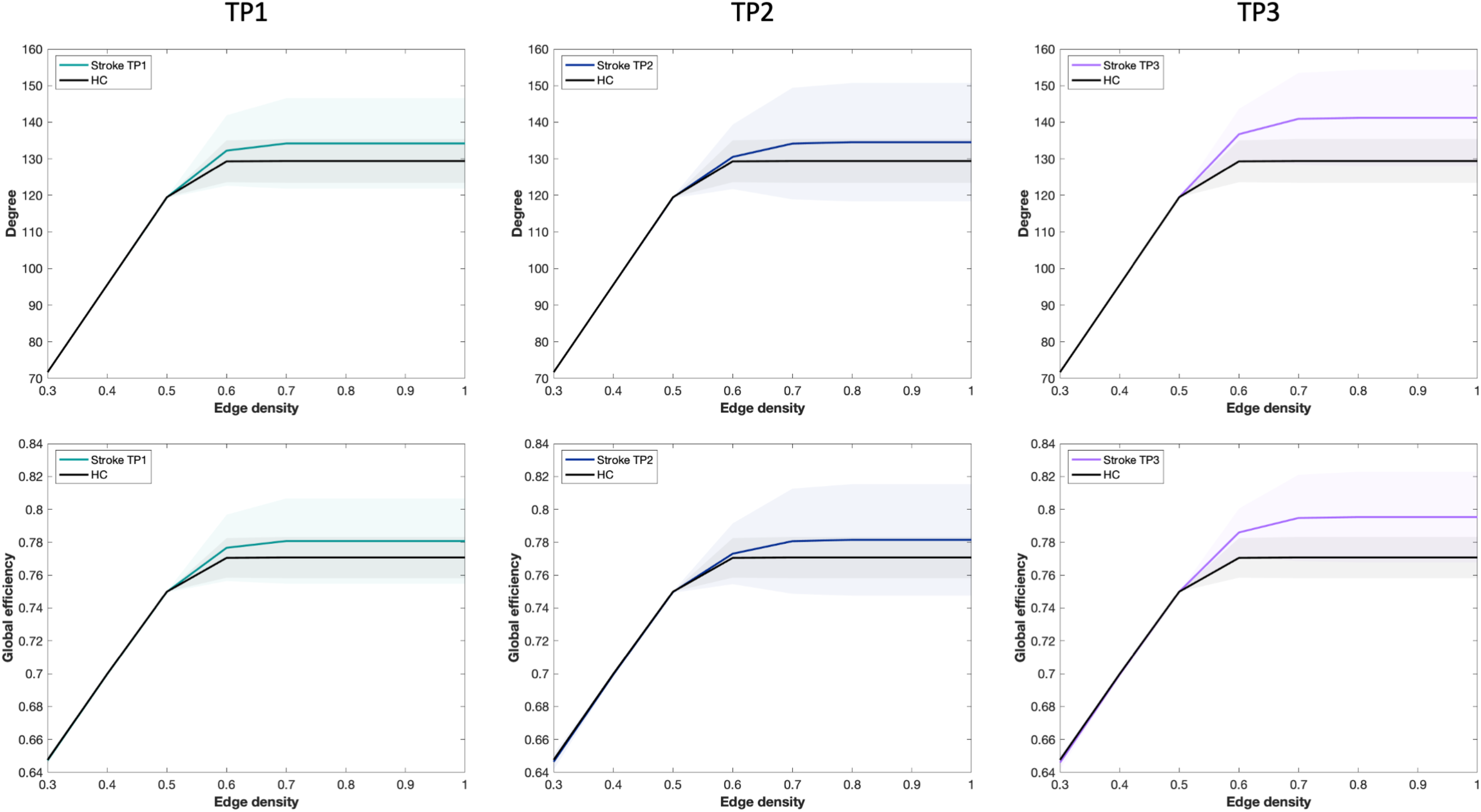
Comparison of network cost and global efficiency between patients and healthy controls at different timepoints. Representation of mean degree and global efficiency, illustrated with means (lines) and standard deviations (filled areas) in patients and healthy controls (HC) at each time point.

#### 2. Impact of lesion simulations on network resilience

**Random failure (Figure 5)**. There were significant changes in E_glob_ over time after random attacks (mixed model, F=4.236, p=0.025). E_glob_ did not differ after random attacks between patients and controls at TP1 (pBH =0.279) nor TP2 (pBH =0.279). However, patients displayed higher E_glob_ at TP3 compared to controls (pBH = 0.012) and to TP2 (pBH = 0.012)).

**Figure 5.**
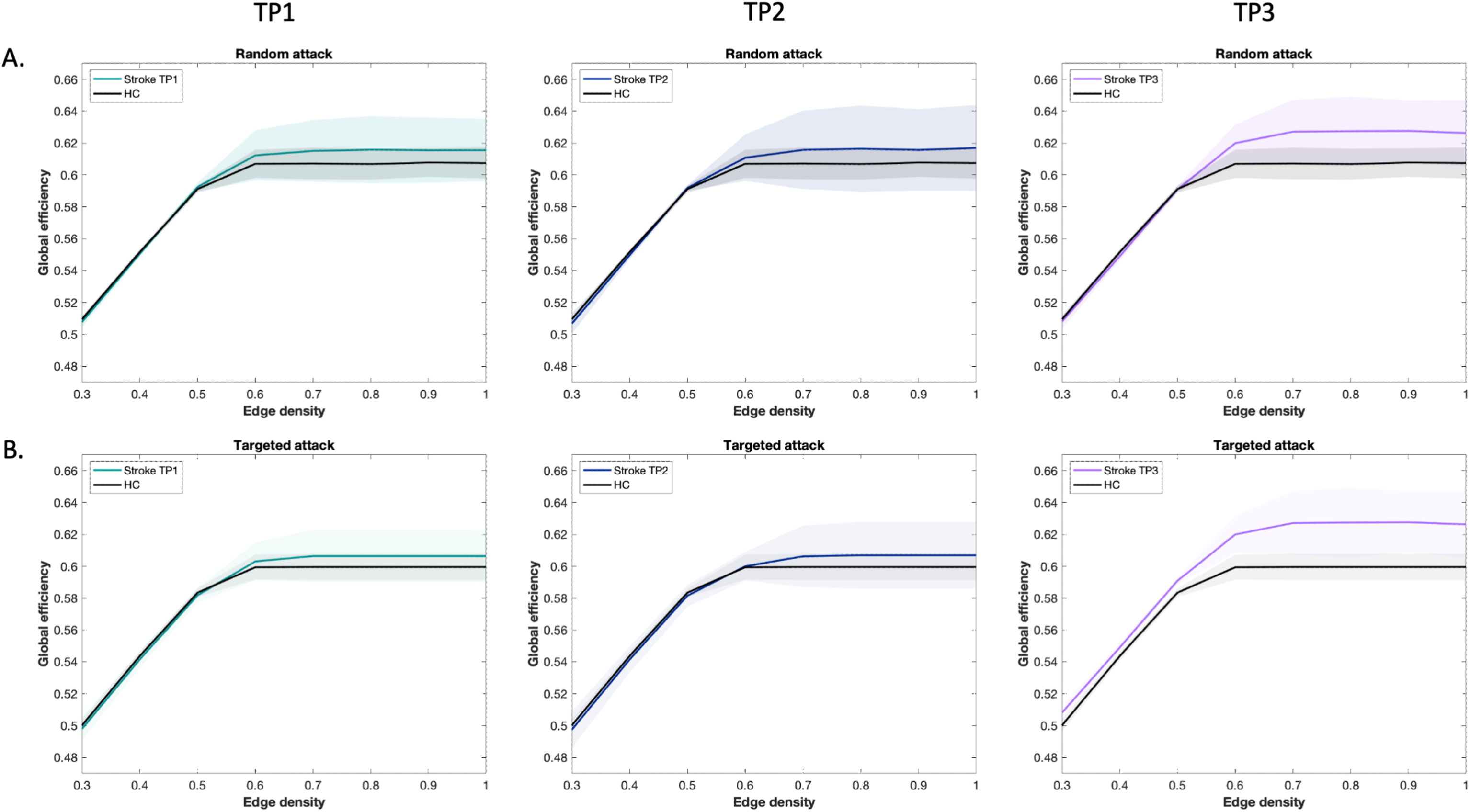
Network resilience after serial random and targeted attacks. Representation of global efficiency after (**A**) Random node deletion and (**B**) Targeted node deletion based on degree. This representation is illustrated with mean global efficiency (lines) and standard deviations (filled areas) after attacks for patients at TP1, TP2 and TP3, as well as for healthy controls (HC).

**Targeted attack**. Using the measure of degree to target serially the nodes of the network, E_glob_ differed from control only at TP3 (pBH = 0.015). Longitudinally, E_glob_ significantly varied across time points (mixed model, F=6,176, p=0.006). E_glob_ increased from TP2 to TP3 (pBH =0.015) but not between TP1 and TP2 (pBH = 0.846).

**Clinically representative lesions. (Figure 6)** E_glob_ varied significantly over time (mixed model, F=16.183, p<0.001). There was no significant change in global efficiency from TP1 to TP2 after all attack types (cortical: p_BH_ = 0.934; subcortical: p_BH_ = 0.916; cortico-subcortical: p_BH_ = 0.788). At TP3 compared to TP2 however, a higher resilience was observed after all types of attacks (cortical: p_BH_ = 0.045; subcortical: p_BH_ = 0.045; cortico-subcortical: p_BH_ = 0.047). When patients were compared to controls, a higher resilience was only found at TP3 (cortical: p_BH_ = 0.012; subcortical: pBH =0.009; cortico-subcortical pBH =0.009) but not at TP1 (cortical: p_BH_ = 0.261; subcortical: p_BH_ = 0.261; cortico-subcortical: p_BH_ = 0.245) nor at TP2 (p_BH_ = 0.261 in all cases).

**Figure 6.**
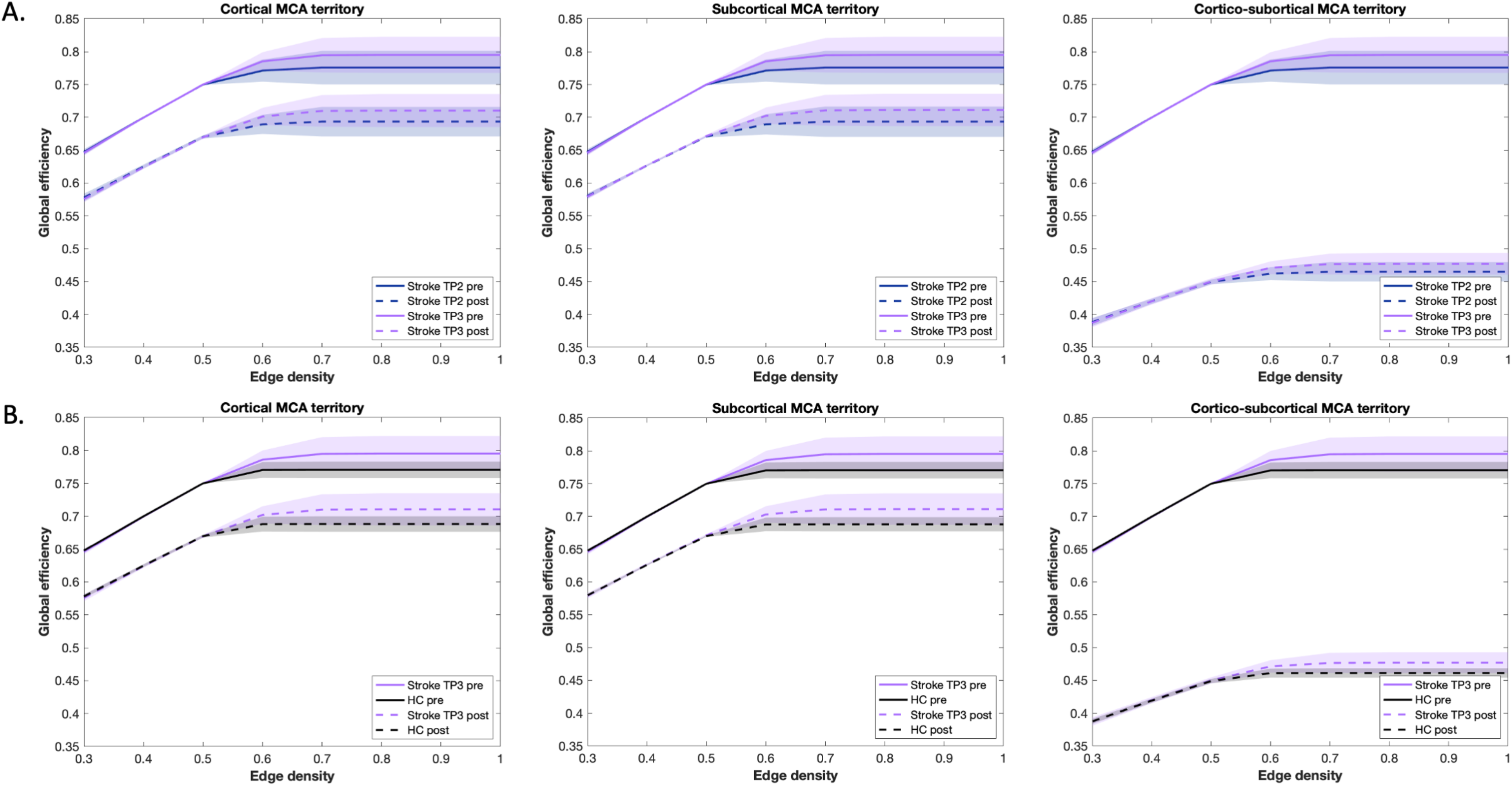
Network resilience after simulation of clinically representative lesions. Global efficiency after cortical, subcortical, and cortico-subcortical simulation of middle cerebral artery (MCA) strokes. (**A**) Pre- and post-lesion comparison in patients at TP2 and TP3. (**B**) cross-sectional comparison between healthy controls (HC) and stroke patients at TP3, illustrated with mean global efficiency before (straight lines) and after attack (dotted lines), as well as standard deviations (filled areas) for each group.

#### 3. Correlation between wiring cost and network resilience

Increase in resilience between 3 weeks (timepoint 2) and 3 months (timepoint 3) after stroke was significantly correlated with the increase in wiring cost of the network (Spearman’s rho=0.785; p=0.001; Figure 7).

**Figure 7.**
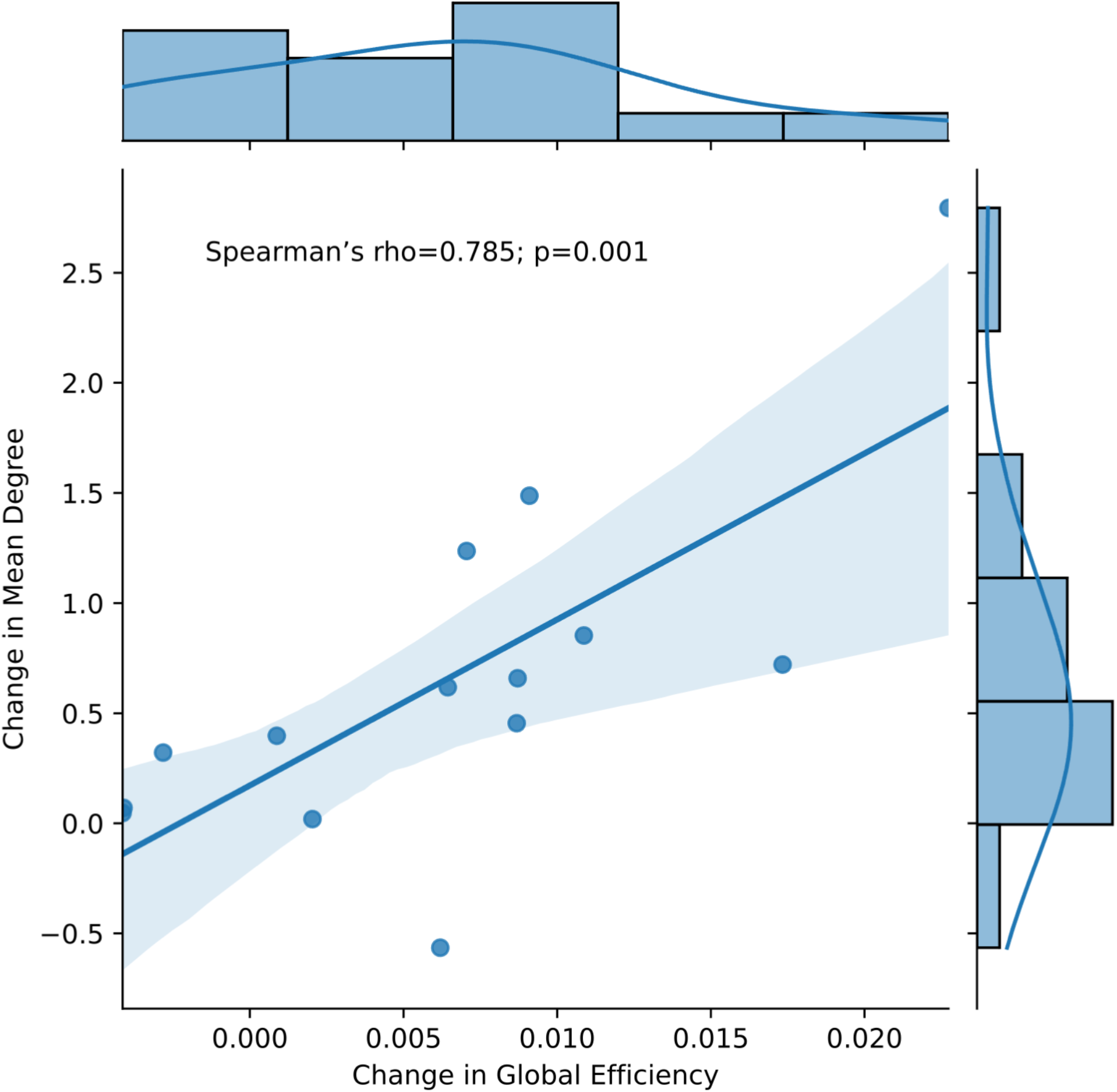
Correlation of change in global efficiency and mean degree between timepoint 2 and 3. Individual values are represented by blue dots. A regression line (straight blue line) is plotted with 95% confidence intervals (shaded area). Distributions for both variables are shown as histograms.

## DISCUSSION

In our population of patients with focal cortical strokes restricted to the primary motor cortex, we showed that resilience in brain networks increased between three weeks and 3 months after stroke. This was demonstrated using attacks mimicking clinically representative stroke by targeting specific or random nodes in the whole brain network. This represents the first evidence that network reorganization may prevent the consequences of a second stroke, at the price, however, of a higher wiring cost.

Resilience to attacks against the network increased at three months compared to 3 weeks, while patients had recovered completely from the first event. This was determined by a higher global efficiency (E_glob_) following lesion simulation of all types. Although network efficiency has not been studied in patients after a second lesion, observational studies have revealed distinct patterns of E_glob_ changes during recovery after a first stroke. In a study including patients with mainly subcortical lesions, a progressive decrease in E_glob_ in the contralateral hemisphere was observed.^24^ When E_glob_ was considered after normalization by random graphs, no changes were described up to 6 months after cortical or subcortical stroke.^25, 26^ More in line with our results, a higher E_glob_ was described within the contralateral hemisphere in mice recovering well from intraluminal occlusion of the right middle cerebral artery.^27^ Comparison of E_glob_ across studies proved however to be challenging, due to the differences in patient population and in the methods applied to determine connectivity (structural vs functional). Importantly, the increase in E_glob_ was obtained at a higher network cost, estimated by mean node degree. The price of shorter paths and of more efficient information propagation and exchanges during stroke recovery could be therefore related to the development of new connections.

We interpret the higher E_glob_ following lesion stimulation at three months as the reflect of a higher capacity of the network to preserve, if a second event occurs, the information integration across the entire network. Because clinical testing was impossible to perform after the lesion simulation, the clinical relevance of these findings remains hypothetical. However, animal studies demonstrated that the longer the event occurs after stroke, the lower the behavioural impact of the recurrent stroke.^2^ Our results may also suggest that the aim of network reconfiguration after stroke may not be limited to the restoration of lost function. New forms of network organization may increase brain resilience to new attacks. If confirmed in a larger stroke population, the results of our study may be relevant to inform neuromodulation strategies that intend to reconfigure network architecture during stroke recovery.

Methodologically, this study opens new perspectives for the study of network reorganisation and resilience in stroke and other diseases. One experimental lesion simulation approach combines MRI-guided brain stimulation with functional connectivity MRI and high-density electroencephalography (EEG). In recent years, brain stimulation, when guided with MRI, has dramatically increased its spatial precision and high-density EEG provides a detailed measure of brain physiology. ^28, 29^ If this strategy represents one of the most promising technique to study brain resilience, it does not allow to mimic lesions that precisely correspond to those commonly observed in acute stroke patients. The use of node deletion to simulate lesions combines several essential characteristics for the study of brain resilience in human. First, it is a highly controllable and precise intervention, that is fully non-invasive. So far, studies using node deletion to simulate lesions, have been used to test the architecture of healthy or pathological networks with no intention to mimic clinically representative strokes.^9, 20, 30-34^. Here, we simulated clinically representative lesions by deleting contiguous nodes using masks of lesions observed in patients admitted to our stroke center. MCA lesions of various sizes and locations were considered in this proof of concept study. Large cortico-subcortical MCA lesions had a more dramatic effect on efficiency than lesions limited to the deep perforator of the MCA (subcortical lesion) or restricted to a superficial branch of the MCA (cortical lesion).

Lesion simulation is an innovative approach, which may have major implications for stroke therapy. In stroke patients, individualized neuromodulation strategies could be developed using TMS or tDCS to not only improve clinical function but also promote resilient network reconfigurations to limit the consequences of future attacks. This approach could also have important implications beyond the stroke field to support the development of individualized therapies for instance in neurodegenerative diseases, such as Alzheimer disease. Identification and promotion of network configurations that are more resilient to the degenerative process may have a huge clinical impact.^35^ Network resilience has also been evaluated in patients at risk to develop post-traumatic stress disorder (PTSD) to tailor preventive cognitive therapies.^36^

There are several limitations to this study. First, we included only patients with discrete lesions limited to the primary motor cortex, who fully recovered at 3 weeks. In patients with larger lesions in the MCA territory, a more severe impact on brain network topology could be expected.^37^ One of the advantages of our study population is the homogeneity of the lesions in term of size and location. Further, because patients have fully recovered clinically at 3 weeks, we were able to make the assumption that changes in connectivity occurring later were consistent with changes in resilience and were not or only partially related to motor recovery.^15, 38-41^ Second, we arbitrarily simulated three cortical and subcortical lesions according to representative ischemic lesions. Stroke in vascular territories including highly central hubs may have a greater impact on efficiency.^11^ Further studies may investigate these hypotheses.

## CONCLUSION

Our results suggest that the optimal network reconfiguration following stroke may not be identical to pre-stroke architecture. Natural selection may increase the robustness of neural networks by favoring their adaptability to unforeseen events.^42, 43^ Clinically, our results indicate that topological resilience may confer protection against the consequence of a novel stroke. In the future, simulation of clinically relevant lesions may have major implications for stroke therapy but also for other neurological and psychiatric diseases. Individualized neuromodulation strategies could be developed to improve clinical function but also promote a resilient network architecture to limit the consequences of future attacks.

## Source of Founding

This work was supported by the Swiss National Science Foundation (320030_166535), the Privat Kredit Bank (PKB) Foundation, and the de Reuter Foundation.

## ABBREVATIONS

AUC: Area Under the Curve
Eglob: Global Efficiency
FDR: False Discovery Rate
M1: Primary motor cortex
MCA: Middle Cerebral Artery
TMS: Transcranial Magnetic Stimulation
TP: Time Point

